# AdipoQ – a simple, open-source software to quantify adipocyte morphology and function in tissues and *in vitro*

**DOI:** 10.1101/2022.06.04.494793

**Authors:** Katharina Sieckmann, Nora Winnerling, Mylene Huebecker, Philipp Leyendecker, Dalila Ribeiro, Thorsten Gnad, Alexander Pfeifer, Dagmar Wachten, Jan N. Hansen

## Abstract

The different adipose tissues can be distinguished according to their function. For example, white adipose tissue (WAT) stores energy in form of lipids, whereas brown adipose tissue (BAT) dissipates energy in the form of heat. These functional differences are represented in the respective adipocyte morphology: whereas white adipocytes contain large, unilocular lipid droplets, brown adipocytes contain smaller, multilocular lipid droplets. However, an automated, image-analysis pipeline to comprehensively analyze adipocytes *in vitro* in cell culture as well as *ex vivo* in tissue sections is missing. We here present AdipoQ, an open-source software implemented as ImageJ plugins that allows to analyze adipocytes in tissue sections and *in vitro* after histological and/or immunofluorescent labelling. AdipoQ is compatible with different imaging modalities and staining methods, allows batch processing of large datasets and simple post-hoc analysis, provides a broad band of parameters, and allows combining multiple fluorescent read-outs. Thereby, AdipoQ is of immediate use not only for basic research but also for clinical diagnosis.

## Introduction

The adipose tissue (AT) is one of the largest endocrine organs in the body and is key in lipid storage and the release of energy (Scherer, 2006; Villarroya *et al*., 2017). The AT is organized according to function. For example, white adipose tissue (WAT) maintains systemic energy balance through the storage and release of free fatty acids and via the secretion of adipokines, whereas the brown adipose tissue (BAT) dissipates energy in the form of heat (Rosen and Spiegelman, 2014; Pfeifer and Hoffmann, 2015; Shamsi *et al*., 2021). The different functions of brown vs. white adipocytes are mirrored in their different morphological appearance: white adipocytes contain large, unilocular lipid droplets, whereas brown adipocytes contain smaller, multilocular lipid droplets (Rosen and Spiegelman, 2014). Furthermore, depending on environmental stimuli, WAT may also contain beige adipocytes that resemble function and appearance of brown adipocytes (Rosen and Spiegelman, 2014). The AT is a dynamic tissue that responds to environmental stimuli and changes its size accordingly. In turn, the contribution of the AT to body weight can vary. Feeding studies in rodents illustrate the fast dynamics of the AT: Within only one week of high-fat feeding, rodent adipocytes become enlarged and can store a multiple-fold of triglycerides per cell compared with baseline cells. Furthermore, the visceral adipose tissue can double within one week if high-fat feeding is initiated (Kleemann *et al*., 2010). The AT expands in two different ways: 1. by increasing adipocyte numbers (hyperplasia) or 2. by increasing adipocyte size (hypertrophy) (Haczeyni *et al*., 2018; Ghaben and Scherer, 2019). How AT expands is a critical determinant for the metabolic homeostasis: hyperplasia has been associated with a metabolically healthy obese state, whereas hypertrophy is strongly connected to the development of obesity-related pathologies, such as metaflammation (a low-grade, chronic inflammation), insulin-resistance, type II diabetes, and arteriosclerosis (Vegiopoulos *et al*., 2017; Ye *et al*., 2021). Obesity-related metabolic disorders like cardiovascular disease are among the most prevalent causes of death worldwide as obesity has reached epidemic proportions in most of the Western world (O’Neill and O’Driscoll, 2015; Hotamisligil, 2017). Thus, understanding the molecular mechanisms underlying AT expansion, in particular hyperplasia vs. hypertrophy, is key to develop novel concepts to target obesity and the related metabolic disorders. To this end, a precise analysis of adipocyte morphology and function, both *in vitro* in tissue culture and in different tissues *ex vivo*, is crucial to shed light on the underlying principles.

So far, automated image analysis to quantify adipogenesis *in vitro* only allowed to determine lipid droplet accumulation (Adomshick *et al*., 2020). For analysing stained tissue sections, several approaches have been described that allow to determine adipocyte count and size distribution (Chen and Farese, 2002; Berry *et al*., 2014; Parlee *et al*., 2014; Maguire *et al*., 2020; Hu *et al*., 2021). Yet, these approaches are only partially automated (Chen and Farese, 2002; Parlee *et al*., 2014; Maguire *et al*., 2020; Hu *et al*., 2021), rely on commercial software (Chen and Farese, 2002; Parlee *et al*., 2014), and/or involve manual, user-biased threshold steps (Parlee *et al*., 2014; Hu *et al*., 2021), limiting throughput, reproducibility, and comparability. Of note, one approach is no longer available (Berry *et al*., 2014).

Here, we present AdipoQ, a set of two open-source plugins for the freely available image analysis software ImageJ (Schneider et al., 2012; Rueden et al., 2017) and its extended version FIJI (Schindelin et al., 2012). AdipoQ provides a simple, versatile, and fully automated analysis of adipocytes in tissue sections and *in vitro* after histological and/or immunofluorescent labelling. AdipoQ allows batch processing of large datasets, simple post-hoc analysis, provides a broad band of parameters, and allows combining multiple fluorescent read-outs.

## Results & Discussion

### AdipoQ

AdipoQ analyzes images acquired from histological or immunofluorescence stainings. It constitutes a two-step workflow consisting of the two ImageJ plugins AdipoQ Preparator and AdipoQ Analyzer (Figure 1A). The AdipoQ Preparator preprocesses the image by segmenting it into fore- and background and creating a mask. The AdipoQ Preparator allows to select from different segmentation strategies that include a combination of image preprocessing and intensity-thresholds or machine-learning-based predictions with StarDist (Schmidt *et al*., 2018b). The latter only applies if a suitable, pre-trained model is available. We provide quick-start user guides and examples to allow the user to quickly find suitable AdipoQ Preparator settings for their data set. AdipoQ Analyzer quantifies the produced mask and outputs various parameters: the number of individual objects (i.e., adipocytes, lipid droplets, or nuclei), the size of each object, the intensity of each object in all image channels, and the intensity surrounding the object in all image channels (Supplementary Table 1). The output files from the AdipoQ Analyzer can be readily read-in into excel or R for further post-hoc analysis. We also provide an R script to merge data from different files.

**Figure 1:**
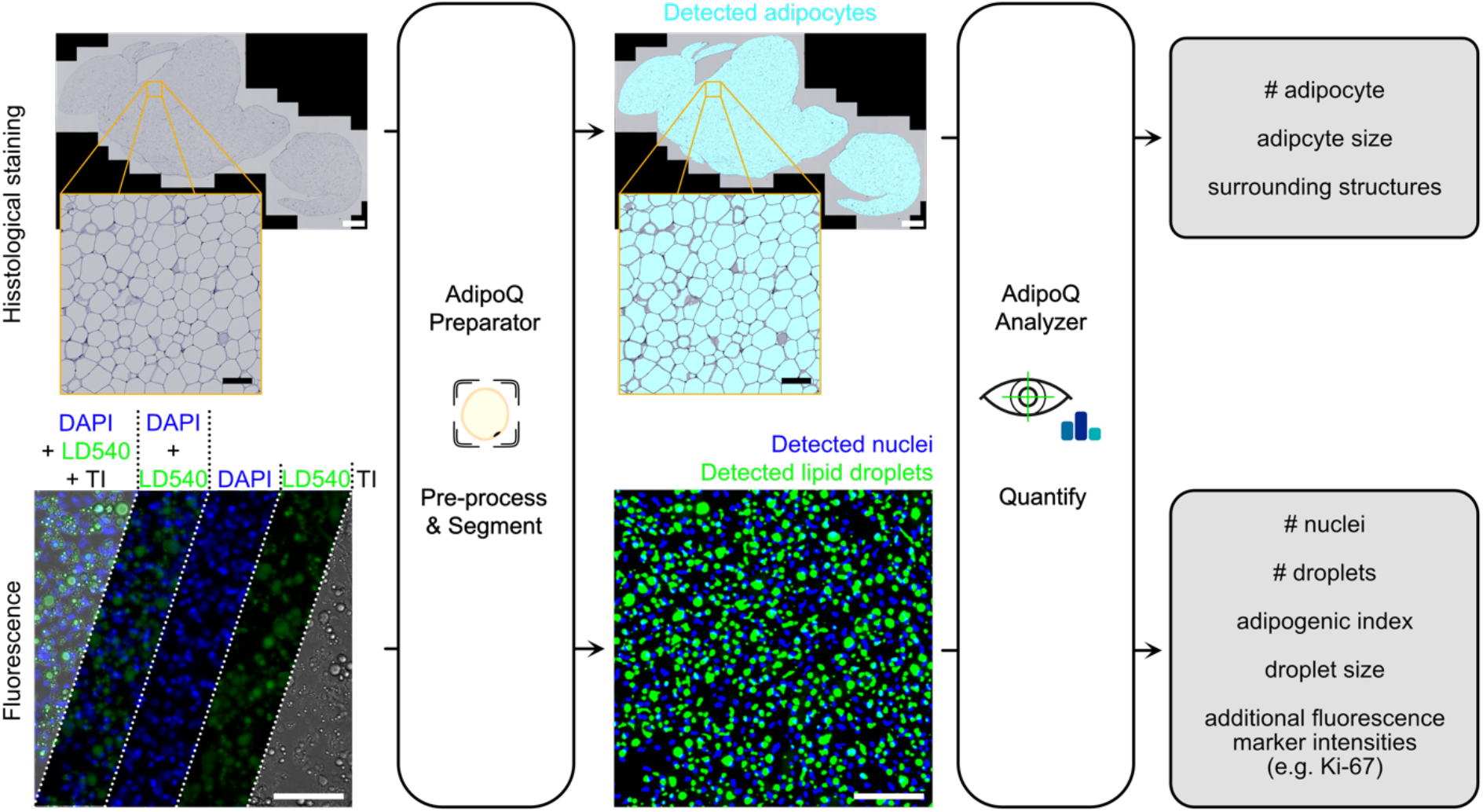
AdipoQ workflow. AdipoQ constitutes a two-step workflow based on the two ImageJ plugins, AdipoQ Preparator and AdipoQ Analyzer, and is applicable to multi-channel images from histological stainings (example image: RGB image) and fluorescence microscopy (example image: epifluorescence microscopy image of cells labeled with DAPI to label nuclei, LD540 to label lipid droplets; TI: transillumination image visualizing the cell structure). The AdipoQ Preparator pre-processes the images for optimal segmentation and subsequently segments them into fore- and background, generating a mask that reveals the detected structures (i.e., adipocytes, nuclei, droplets). The AdipoQ Analyzer quantifies this mask. AdipoQ Analyzer counts the structures and determines the size of each structure and the intensity levels within and adjacent to each structure, in all different channels in the image. This allows to measure also additional fluorescent markers. Scale bars in top row: 1 mm (Inset: 100 µm). Scale bars in bottom row: 100 µm.

AdipoQ provides transparent analysis methods and is freely accessible. The source code, the plugins, and a comprehensive user guide are freely available through the GitHub repository https://github.com/hansenjn/AdipoQ. Being integrated in the broad and free analysis software ImageJ, the AdipoQ plugins can be controlled via a user interface and do not require coding knowledge. Furthermore, the plugins contain a batch processing tool that allows to automatically process a list of files consecutively without user interaction. Using the AdipoQ Preparator in FIJI, raw microscopy formats can also be directly loaded. Thus, we present a very straight-forward and fully automatized workflow that allows to analyze a dataset from raw data to plots in only three fully automated steps: two ImageJ plugins and automated post-hoc analysis with an R script. In contrast to previous approaches (Chen and Farese, 2002; Berry *et al*., 2014; Parlee *et al*., 2014; Maguire *et al*., 2020; Hu *et al*., 2021), AdipoQ, with its batch-processing modalities and easy accessibility through ImageJ and FIJI provides high-throughput analysis, freely available for all users. Furthermore, AdipoQ determines a broader set of parameters and also allows to combine read-outs from multiple fluorescence channels.

### Analyzing AT sections *ex vivo* using AdipoQ

We demonstrate the strength of AdipoQ using histological stainings of WAT sections and fluorescence labeling of different adipocytes cultured *in vitro*. As a benchmark for the analysis of AT, we studied WAT expansion in tissue sections labeled with Hematoxylin-Eosin (HE) from mice that were fed with a high-fat diet (HFD) for 8 weeks compared to mice fed with chow diet (CD) (Figure 2). On HFD, the adipocyte size was visually larger compared to CD (Figure 2A). We applied the AdipoQ analysis pipeline and determined adipocyte size and its distribution (Figure 2, B and C). For both female and male mice, larger adipocytes were significantly more frequent while smaller adipocytes were significantly less frequent in WAT from mice on HFD compared to mice on CD, indicating that the WAT expanded by adipocyte hypertrophy and verifying previous results (Gao *et al*., 2015).

**Figure 2:**
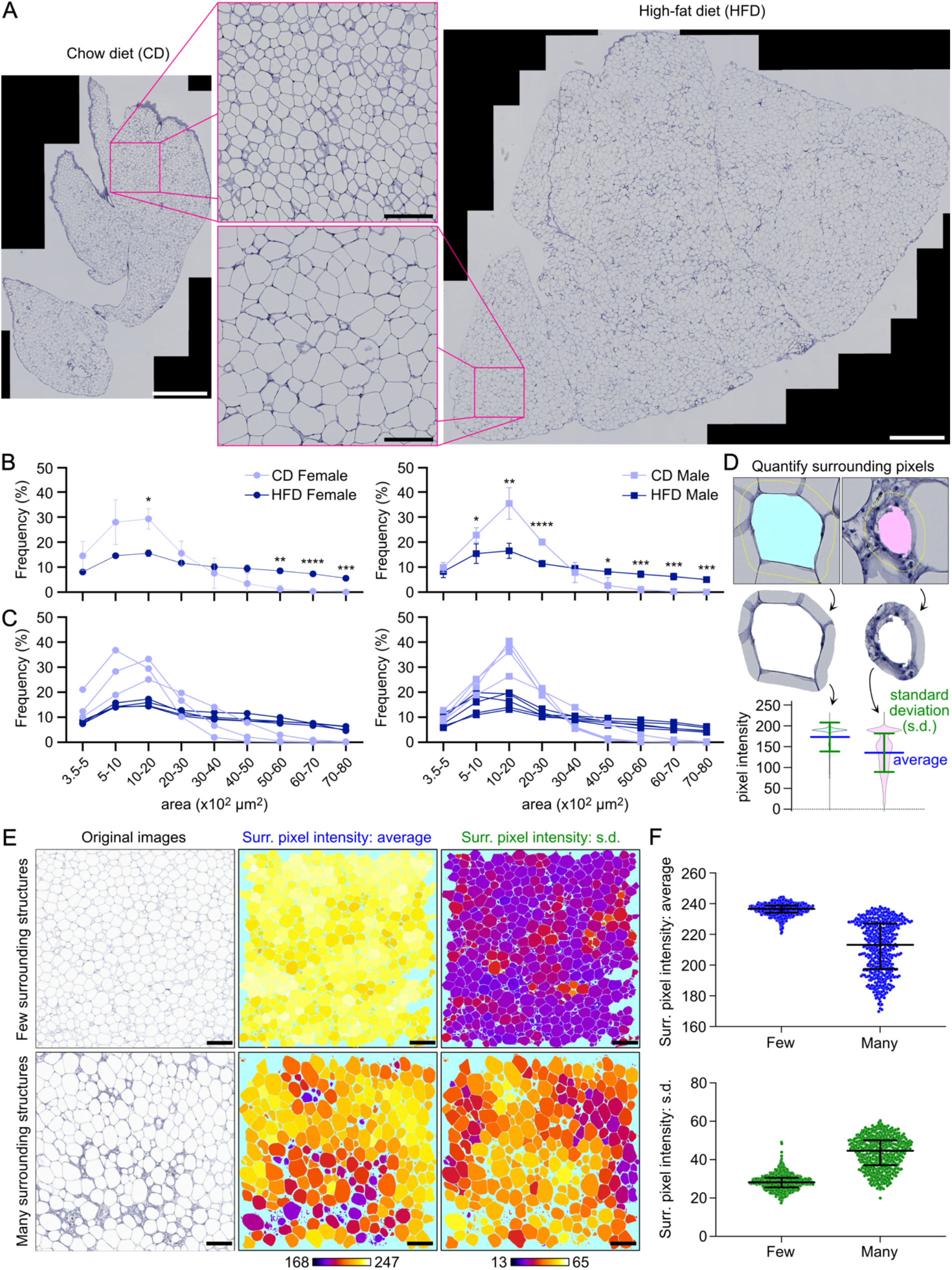
AdipoQ characterizes adipocytes in images of histological stainings. **(A)** Example images of Hematoxylin-Eosin-stained adipose tissue extracted from male wild-type mice fed chow diet (CD, left) or high-fat diet (HDF, right). Scale bars: 1 mm (Inset: 200 µm). **(B-C)** Adipocyte size distribution quantified by analyzing images as exemplified in (A), acquired from female and male wild-type mice on CD (n = 3 (female), n = 4 (male)) or HFD (n = 4 (female), n = 5 (male)). Per mouse, 2-6 sections were quantified. **(B)** Mean ± standard deviation of the distributions from all mice per group. p-values indicated for unpaired, two-sided t-tests with Welch correction between CD and HFD at the size range; *, p ≤ 5×10^−2^; **, p < 10^−2^; ***, p < 10^−3^; ****, p < 10^−4^. If no symbol is indicated, the comparison was non-significant. **(C)** Individual distributions. Each line represents one mouse. **(D-F)** Quantification of the adipocyte surrounding. **(D)** AdipoQ allows to quantify the surrounding of individual adipocytes. It extracts the pixel intensities within a defined radius around the adipocyte and determines, among other parameters (see Supplementary Table 1), the average and standard deviation of the pixel intensities in the surrounding (shown as bars in the violin plots). Such parameters reveal non-adipocyte structures in the surrounding. **(E-F)** AdipoQ analysis of example images from two different mice, showing adipose tissue with few and with many surrounding structures. **(E)** Maps (middle and right, generated by AdipoQ) that visualize the detected adipocytes for the example images (left), color-coded by average (middle) or standard deviation (s.d., right) of surrounding pixel intensities. Scale bar: 200 µm. **(F)** AdipoQ analysis results for the images shown in (E). Data points show individual adipocytes. Bars show median and interquartile range.

Several stainings allow to assess the integrity of the AT and the interaction of adipocytes with other cells, e.g., when labeling collagen as a fibrosis marker (Cinti *et al*., 2005; Harman-Boehm *et al*., 2007; Strissel *et al*., 2007; Murano *et al*., 2008). For examples, in WAT sections from mice on HFD, accumulation of other, non-adipocyte cells was visible around adipocytes in an HE staining (Figure 2D). These so-called crown-like structures are histologic hallmarks of a proinflammatory process in the AT, representing dying adipocytes surrounded by macrophages (Murano *et al*., 2008). AdipoQ features parameters that quantify the surrounding of the analyzed object (i.e., the adipocyte). To this end, the pixel intensities within a user-defined distance are extracted and quantified (Supplementary Table 1, Figure 2D). For example, the standard deviation and average of pixel intensities in the surrounding of the adipocyte are determined and describe the extent of non-adipocyte structures surrounding the adipocyte: Since surrounding structures other than adipocytes are more heterogeneous in intensity and of lower intensity compared to the “bright” adipocytes in the images, the average and standard deviation of surrounding pixel intensities are lower or higher, respectively, the more non-adipocyte structures are present (Figure 2D). To demonstrate the application of these parameters, we extracted two field of views from the analyzed HE-labelled WAT sections, one with only a few and one with many visible non-adipocyte structures and used AdipoQ to determine average and standard deviation of the pixel intensities in the surroundings of the adipocytes (Figure 2, E and F). AdipoQ clearly revealed the differences in non-adipocyte structures between the two images, demonstrating that these two parameters can serve as indicators for the adipocyte microenvironment (Figure 2F). Importantly, it is also possibly to perform a combined HE and IHC staining, e.g., for macrophages (Lee *et al*., 2011). Here, AdipoQ could be used to detect adipocytes in HE labelled tissue sections and reveal macrophage accumulation in the adipocyte surrounding based on the macrophage channel.

To ensure a broad applicability of AdipoQ, we tested AdipoQ on images of HE-labelled tissues from other labs: (1) histological images of human visceral (Supplementary Figure 1A) and subcutaneous (Supplementary Figure 1B) adipose tissue from patients of different sex and age, downloaded from the Genotype-Tissue Expression (GTEx) portal (https://gtexportal.org/) (Supplementary Figure 1, A and B) and (2) images from a freely-available murine adipose tissue data set (Casero *et al*., 2021) (https://dx.doi.org/10.5281/zenodo.5137433), in which cell borders between adipocytes were particularly weak (Supplementary Figure 1C). In all tested images, AdipoQ successfully detected adipocytes under default settings (Supplementary Figure 1, A-C).

### Analyzing adipogenesis *in vitro* using fluorescence labeling and AdipoQ

The regulation of adipocyte progenitor cell (APC) differentiation determines whether the AT expands via hypertrophy or hyperplasia. This process is termed adipogenesis and can be analyzed *in vitro* using primary APCs or immortalized cell lines, e.g., 3T3-L1 cells, which have been generated from a 3T3 mouse fibroblast sub-strain and are committed to the adipocyte lineage (Green and Meuth, 1974). To this end, reliable image analysis tools to assess adipogenesis using fluorescence read-outs are required.

### Analyzing adipogenesis of 3T3-L1 cells *in vitro* using AdipoQ

We acquired images of 3T3-L1 cells before and after induction of adipogenesis (Figure 3A) and analyzed them using AdipoQ (Figure 3, B-C). To visualize lipid droplets, we used an antibody directed against perilipin (PLIN1), a protein that associates with the lipid droplet membrane (Brasaemle *et al*., 2009). To visualize nuclei, we co-stained the cells with DAPI. Using AdipoQ, we quantified lipid droplet accumulation by determining the PLIN1^+^ area and defined an adipogenic index by calculating the ratio of the PLIN1^+^ area and the DAPI^+^ area (Figure 3B). Lipid accumulation was already visible after 2 days of induction and increased over the 8 day time course of differentiation (Figure 3, A and B). In untreated control cells, lipid accumulation was much lower as indicated by a lower adipogenic index compared to fully differentiated cells (Figure 3, A and B). Thus, the AdipoQ output parameters allow to reliably quantify adipogenesis with high throughput in an automated fashion.

**Figure 3:**
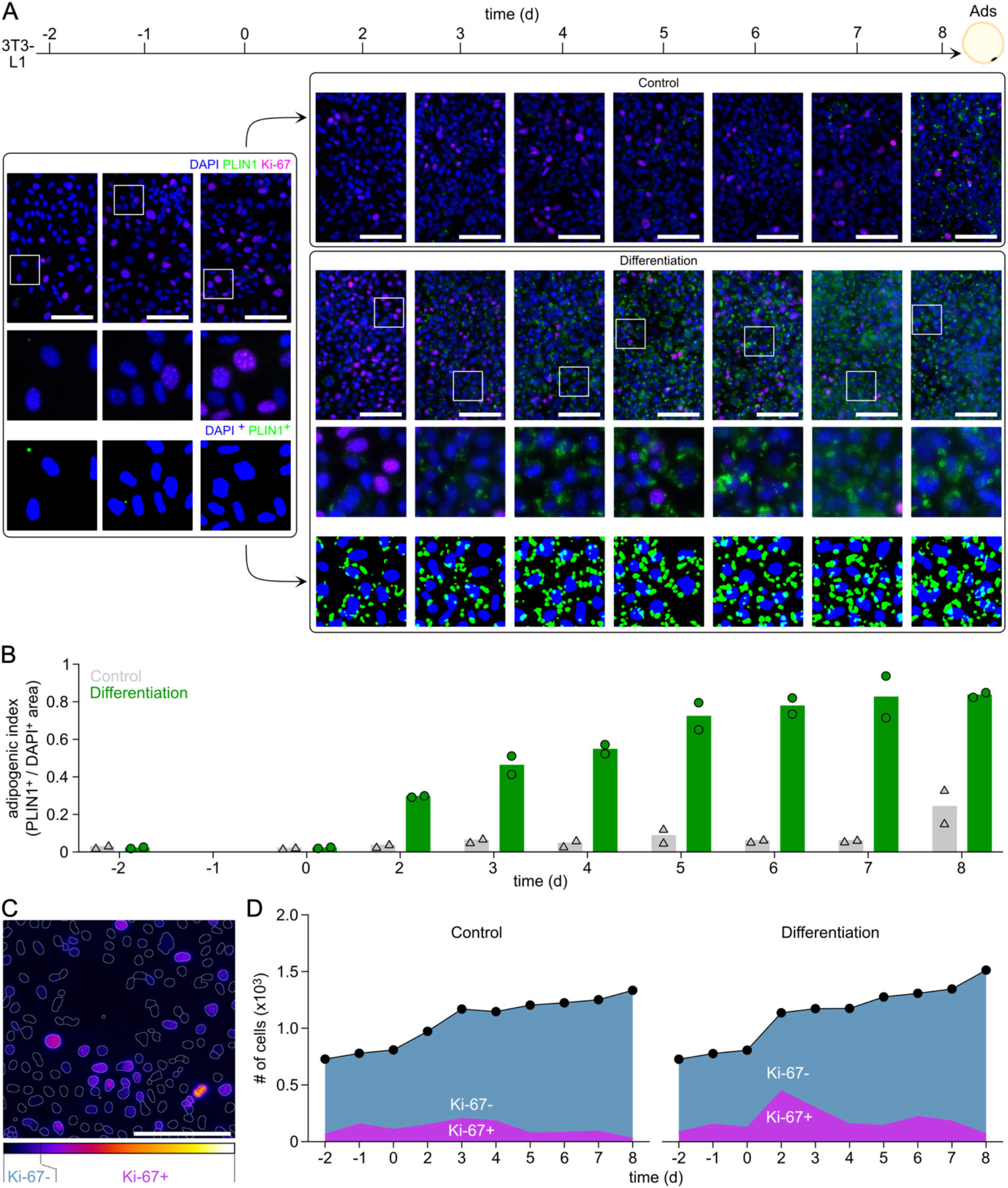
AdipoQ quantifies adipogenesis and proliferation *in vitro*. **(A)** Example images of 3T3-L1 cells during adipogenic differentiation, which was induced at day 0. Cells were fixed at indicated time points. Cells were stained with Ki-67 (magenta), PLIN1 (green), and DAPI (blue). Scale bar = 100 µm. Rectangles indicate magnified views below, for which also the detected DAPI^+^ and PLIN1^+^ areas are indicated. **(B-D)** AdipoQ analysis of the imaging data presented in (A) (n = 1 example experiment). **(B)** Adipogenic index, determined by calculating the total lipid droplet area (PLIN1^+^) and dividing it by the total nuclei area (DAPI^+^). Bars show mean, data points show individual experiments (simultaneously performed, average of four images per experiment). **(C)** Quantifying adipocyte proliferation using Ki67 staining. Ki67 channel for an example image shown (colored by the look-up table indicated below). Masks for individual nuclei (overlayed with white lines in image) are determined from the DAPI channel. For each nucleus mask, the Ki67 signal is determined and if the median intensity within the mask exceeds a fixed threshold, the nucleus is considered Ki67^+^. **(D)** Proliferation kinetics of 3T3-L1 cells as indicated by the total number of nuclei and the nuclei that showed a high Ki-67 intensity (Ki-67^+^). Data pooled from both experiments.

Accumulation of lipid droplets is the most intuitive feature of adipocyte differentiation; however, many other molecular processes are differentially regulated during adipogenesis. AdipoQ allows to measure additional fluorescence markers to follow the kinetics of any protein of interest during differentiation. We tested this by additionally labeling Ki-67, a protein whose presence is closely linked to the cell cycle und, thus, serves as a proliferation marker (Brown and Gatter, 2002; Li *et al*., 2015). As Ki-67 localizes to the nucleus, we measured the signaling intensity of Ki-67 in the nuclei (Figure 3C). To this end, we detected individual, DAPI^+^ nuclei with AdipoQ. By combining image segmentation with ImageJ’s Watershed method or, alternatively, using a machine-learning-based nuclei detection, AdipoQ allows to identify single nuclei and in turn, reveal 1) the number of nuclei per image and 2) the pixel intensities within each nucleus for any other fluorescence channel (i.e., the Ki-67 signal per nucleus). AdipoQ outputs different pixel-intensity parameters, e.g., average intensity, median intensity, minimum or maximum intensity (Supplementary Table 1). To identify Ki-67^+^ nuclei, the median intensity parameter was chosen, as this is least sensitive to noise. To distinguish Ki-67^-^ from Ki–67^+^ nuclei, a fixed threshold was set based on the intensity histogram. The ratio of Ki-67^+^ nuclei visualized the proliferation kinetics during adipogenesis, revealing clear differences between control cells and cells induced to differentiate (Figure 3D). Proliferation increased after 2 days of adipogenesis induction and decreased with increasing lipid accumulation (Figure 3, A and D). In contrast, proliferation in untreated control 3T3-L1s was generally lower (especially at day 2) and decreased over time (Figure 3D).

### Analyzing adipogenesis of primary APCs *in vitro* using AdipoQ

To test if our AdipoQ analysis pipeline can also be applied to primary cells, we generated images from primary murine and human APCs and analyzed them using AdipoQ. We isolated APCs from WAT, BAT, and bone marrow (BM) of wild-type mice and from human BAT from the supraclavicular area, cultured them *in vitro*, and induced adipogenesis. Here, we visualized lipid droplets using the lipophilic fluorescence dye LD540 (Spandl *et al*., 2009) and quantified lipid accumulation by measuring the LD540^+^ area in a field of view using AdipoQ. Similar to the 3T3-L1 cells, concomitant staining with DAPI allowed to quantify the adipogenic index as ratio of the LD540^+^/DAPI^+^ area. Lipids accumulated in WAT-APCs after inducing adipogenesis over 7 days compared to untreated controls (Figure 4A), similarly to adipogenesis in 3T3-L1 cells (Figure 3, A and B). After 7 days, lipid accumulation was also observed in untreated control cells, but to a much lesser extent. Similar results were observed for APCs from murine BAT and BM (Figure 4, B and C) and from human BAT (Supplementary Figure 2, A and B).

**Figure 4:**
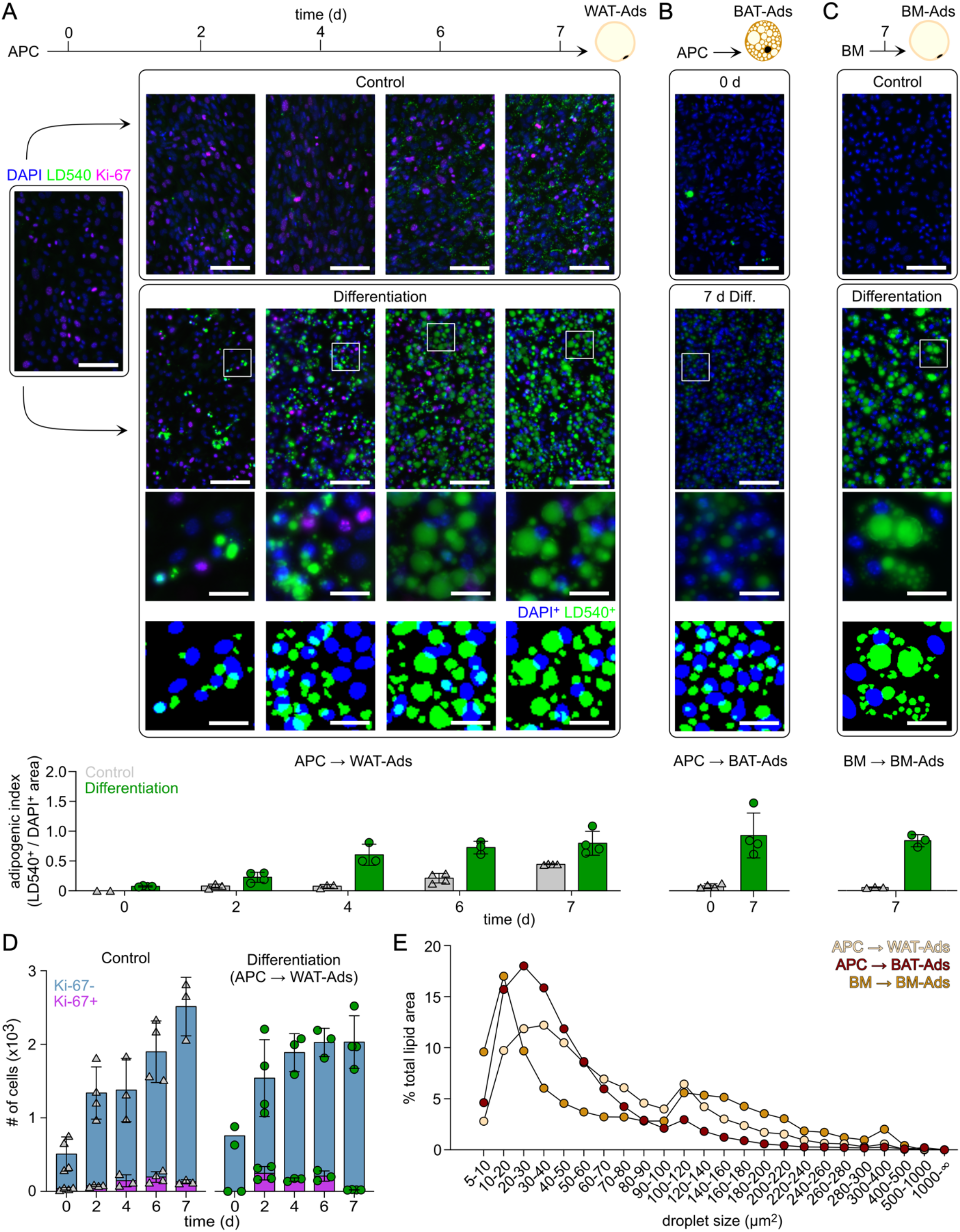
Differences in adipogenesis parameters in primary APC models can be assessed with AdipoQ. AdipoQ analysis of APCs isolated from WAT, BAT, and BM before and after differentiation. **(A)** Example images (top) and adipogenic index determined by AdipoQ (bottom) for APCs isolated from WAT. Rectangles indicate magnified views below, for which also the detected DAPI^+^ and LD540^+^ areas are indicated. Scale bars = 100 µm (magnified views: 25 µm). To visualize proliferation during the course of differentiation, APCs were fixed at different days and stained with Ki-67 (magenta), LD540 (green), and DAPI (blue). The Adipogenic index is determined by the ratio of total lipid droplet area (LD540^+^) to total nuclei area (DAPI^+^). Bars indicate mean ± standard deviation. Shown is an example experiment, data points show individual images. **(B)** See (A), for BAT. Cells were fixed after 0 or 7 days in culture and stained with LD540 (green) and DAPI (blue). Data points show individual experiments (n = 4). **(C)** See (B), for BM. Cells were fixed after 7 days in culture. Data points show individual experiments (n = 3). **(D)** Proliferation kinetics of WAT-APCs determined by the total number of nuclei and the fraction of nuclei that showed a high Ki-67 intensity (Ki-67^+^). Shown is an example experiment, data points show individual analyzed images. **(E)** Quantification of changes in lipid droplet size is depicted as fraction of total lipid area per lipid droplet size. Per condition, lipid droplets were pooled from all experiments and images, from day 7 after differentiation.

Also here, we analyzed proliferation using Ki-67 labeling (Figure 4D): WAT-APCs increased their proliferation rate after 2 days of induction and decreased the rate with increasing lipid storage (Figure 4, A and D). In contrast, proliferation in control WAT-APCs steadily increased and peaked at day 6 (Figure 4D). This is also represented in the total number of nuclei, as control APCs have a higher number of nuclei after 7 days of culture, compared to the differentiated cells (Figure 4D).

AdipoQ also allows to distinguish individual lipid droplets (Supplementary Figure 2A). Accordingly, we determined the lipid droplet size in mature adipocytes (Ads), generated from murine WAT-, BAT-, or BM-APCs *in vitro* (Figure 4E). Assessing lipid droplet size *in vitro* may not be directly translatable to an *in vivo* situation, as two-dimensional cell culture leads to the formation of small, multilocular droplets instead of one solitary droplet (Dufau *et al*., 2021). Nonetheless, we observed differences in multilocular lipid droplet size between the three adipocyte types (Figure 4E): BAT-Ads contained a higher number of smaller lipid droplets compared to WAT-Ads, which contained more lipid droplets in the intermediate and higher size ranges (Figure 4E). BM-Ads contained a high number of small and big droplets but less in the intermediate size range. This highlights that the AdipoQ analysis pipeline allows to identify functional differences between cell types.

### Reproducibility of AdipoQ analysis between different experimental replicates and users

We next scrutinized the reproducibility of the AdipoQ analysis. First, we tested how reproducible the obtained size distributions are across different experimental replicates. We compared the size distributions derived from nine individual human BAT-APC-derived adipocyte cultures, cultured and processed at the same time. The standard deviation of the size distribution was low and ranged from 0.02 (for droplet areas >= 180 and < 200 µm^2^) to 1.35 percent of the total lipid droplet area of all size groups (for droplet areas >= 5 and < 10 µm^2^) (Supplementary Figure 2C).

Next, we aimed to illustrate differences between the analysis results obtained by two different investigators. We compared results for the same images, analyzed by two different investigators who selected different AdipoQ segmentation settings. Whereas the absolute adipogenic index values were different for the two settings, relative changes between control and differentiation medium were similar and both settings detected a clear difference between control and differentiation medium (Supplementary Figure 2D). Similar results were obtained for the size distribution of detected lipid droplets, which also showed only small deviations between both settings (Supplementary Figure 2E). These results demonstrate that for a direct comparison of absolute values, AdipoQ needs to be run using the same settings, whereas relative values and the lipid droplet size distribution are only marginally affected by different settings.

### Accuracy of lipid droplet and nuclei detection by AdipoQ

As shown above, AdipoQ can reliably detect individual adipocytes, droplets, or nuclei in diverse images (Figure 2-4, Supplementary Figure 1-2). The accuracy, however, depends on the quality of sample preparation and of the acquired images. For a correct object detection, object borders need to be of sufficient contrast and well-resolved. Even though AdipoQ can detect also weak borders (Supplementary Figure 1C), it cannot separate adjacent objects if the border is invisible in the image or if the image resolution does not allow to resolve the border clearly. This becomes particularly apparent in the detection of small lipid droplets. We determined the number of falsely detected droplets by AdipoQ and distinguished between multiple droplets detected as one droplet (fused), one droplet detected as multiple droplets (fragmented), and droplets that were not detected at all (missing). Errors for fragmented or missing droplets were low (Supplementary Figure 2, F and G), whereas fused lipid droplet errors occurred frequently, reaching even 36% of all detected objects for one images of BAT-Ads (Supplementary Figure 2H). However, areas of fused objects were relatively small (Supplementary Figure 2I), compared to the overall lipid droplet size distribution (Figure 4E), indicating that they arise from small droplets. Importantly, images acquired at 0.9 µm/px showed a two-fold higher error rate for fused droplets compared to images acquired at 0.45 µm/px (Supplementary Figure 2H), highlighting that fused droplet errors can be reduced by acquiring images at higher magnification/image resolution. Of note, errors from fused or fragmented droplets do not affect global image parameters, such as the adipogenic index, since the index does not count nuclei or droplet but only determines the nuclei and droplet positive areas.

The accuracy of nuclei detection depended mostly on the culture condition and cell type. We observed that the accuracies for nuclei detection was varying across different adipocyte types, with 0.6% for WAT-Ads and 2.6% for BAT-Ads (Supplementary Figure 2J). We noticed that a main source of error in nuclei detection is a too high cell density in 2D culture: In human BAT-Ads, we observed that cells formed small islands with high cell density under control conditions (See upper left in image shown in Supplementary Figure 2A), which resulted in less accuracy of nuclei detection (Supplementary Figure 2K).

### AdipoQ outperforms current alternatives for lipid droplet detection

The only alternative software to quantify individual lipid droplets *in vitro* is a CellProfiler script (Adomshick *et al*., 2020). We tested the script on some of our images. Nuclei detection was possible only with customization and did not perform well for dense culture regions (Supplementary Figure 2L). Lipid droplet detection was by far less accurate than by AdipoQ, since most droplets were fused to larger objects (Supplementary Figure 2L). Nonetheless, it might be possible to establish a new analysis pipeline in CellProfiler, which would achieve a similar lipid droplet detection as performed by AdipoQ. However, right now, AdipoQ appears to outperform this CellProfiler script.

In summary, we report a new method to comprehensively analyze adipocyte morphology *in vitro* in cell culture as well as *ex vivo* in tissue sections, applicable to human and mouse cells/tissues. By combining different imaging modalities and staining methods (histological and fluorescent stainings) as input for the analysis pipeline, AdipoQ allows to distinguish adipocyte types and AT depots, which is of immediate use not only for basic research but also for clinical diagnosis.

## Materials and Methods

### Cell lines and cell culture

The 3T3-L1 cell clone #27 was kindly provided by Prof. Christoph Thiele, LIMES Institute, University of Bonn, Germany. Cells were maintained in DMEM, supplemented with 1% GlutaMAX™-I (both: Gibco/Life Technologies) and 10% foetal calve serum (FCS, Biochrom) at 37 °C and 5% CO_2_. Cells have been tested for mycoplasma twice a year and were free from mycoplasma.

### APC isolation

Gonadal WAT and the interscapular BAT were surgically removed from mice and processed for adipose precursor cell (APC) enrichment as follows.

WAT was minced and digested with collagenase II in 0.5% BSA (Sigma) in PBS at 37°C with agitation. The digestion was quenched by adding AT buffer (0.5% BSA in PBS). The dissociated cells were passed through a 100-μm filter (Corning) and subjected to centrifugation at 500 × g for 10 min. The resulting supernatant containing mature adipocytes was aspirated, and the pellet, consisting of the stromal vascular fraction (SVF), was resuspended in red blood cell lysis buffer (Biolegend) for 2 min at RT. The reaction was stopped by adding AT buffer and centrifugation at 500 × g for 10 min. Cells were then passed through a 40-μm filter (Corning) and then maintained in DMEM/F12 (1:1), supplemented with 1% GlutaMAX™-I, 1% Penicillin-Streptomycin (all Gibco/Life Technologies), 10% FCS (Biochrom), 33 mM biotin (Sigma), and 17 mM D-pantothenate (Sigma) at 37 °C with 5% CO_2_.

BAT APC isolation was adapted based on a protocol by Schmidt *et al*. (2018a). Minced tissue was digested with collagenase II in serum-free medium (DMEM/Ham’s F12, Gibco/Life Technologies) containing 0.5% BSA, 1% glutamine, 33 mM biotin, 17 mM D-pantothenate, and 1% Penicillin-Streptomycin at 37°C with agitation. After digestion, medium containing 10% FCS was added. Cells were then passed through a 100-µm filter (Corning) and centrifuged at 300 x g for 5 min. The supernatant was aspirated and the pellet containing the APCs was resuspended in 10 mL medium containing 10% FCS. Cells were washed again and subsequently, the cell pellet was resuspended in red blood cell lysis buffer (Biolegend) and incubated for 2 min at RT. Reaction was stopped with medium containing 10% FCS. After centrifugation, cells were resuspended in medium containing 20% FCS and seeded on a 6-cm cell culture dish (Greiner). APCs were maintained at 37°C and 5% CO_2_ until cells reached confluency.

BM-APC isolation was based on an adapted protocol by Huang *et al*. (2015). Briefly, femur and tibia were dissociated, and the surrounding soft tissue was removed. After immersion in ethanol (70 %) for 1 minute, the marrow was flushed out with MEM-alpha (Gibco/Life Technologies), supplemented with 15 % FCS and 1% Penicillin-Streptomycin, into a 10 cm culture dish until the bones became pale. BM-APCs were kept in culture at 37 °C with 5 % CO_2_ until the cells reached 80 % of confluence.

Primary human BAT-Ads were isolated and differentiated as previously described (Jespersen *et al*., 2013; Gnad *et al*., 2020). Cells were maintained in 60-mm culture dishes containing DMEM/F12, 10% FBS, 1% Penicillin/Streptomycin (all from Invitrogen) and 1 nM acidic FGF-1 (ImmunoTools). Cells were incubated at 37° C with 5 % CO2.

### *In vitro* adipogenesis assay

For differentiation, 3T3-L1 cells were seeded on CellCarrier Ultra 96-well plates (PerkinElmer). When cells reached confluency, adipogenesis was induced by switching to induction medium, modified from Hilgendorf *et al*. (2019), containing 0.4 µg/mL insulin (Sigma), 0.1 µM Dexamethasone (Sigma), and 20 µM 3-isobutyl-1-methylxanthine (IBMX; Sigma). After 2 days of induction, medium was exchanged to freshly prepared maintenance medium, containing 1 µg/mL insulin. Afterwards, maintenance medium was changed every other day. Additionally, as a negative control, undifferentiated cells were kept in medium.

For differentiation into WAT-Ads or BM-Ads, isolated WAT APCs or BM APCs were seeded on CellCarrier Ultra 96-well plates. When cells reached confluency, adipogenesis was induced by switching to induction medium, modified from Hilgendorf *et al*. (2019), containing 5 µg/mL insulin, 1 µM Dexamethasone, 100 µM IBMX, and 1 µM rosiglitazone (Sigma). After 3 days of induction, medium was exchanged to freshly prepared maintenance medium, containing 1 µg/mL insulin. Afterwards, maintenance medium was changed every other day. Additionally, as a negative control, undifferentiated cells were kept in medium.

For differentiation to BAT-Ads, isolated BAT APCs were seeded on CellCarrier Ultra 96-well plates. Once cells reached confluency, adipogenesis was induced by switching to induction medium containing 850 nM insulin, 1 µM Dexamethasone, 250 µM IBMX, 1 µM rosiglitazone, 125 µM indomethacin (Sigma), and 1 nM 3,3′,5′-Triiodo-L-thyronine (Sigma). After 2 days of induction, cells were maintained in medium containing 1 µM rosiglitazone and 1 nM 3,3′,5′-Triiodo-L-thyronine until day 7. Medium was exchanged every other day. Additionally, as negative control, undifferentiated cells were kept in medium.

Primary human BAT-Ads were differentiated as previously described (Jespersen *et al*., 2013; Gnad *et al*., 2020). Adipocytes were seeded on CellCarrier Ultra 96-well plates. Two days after cells reached confluency, adipogenesis was induced by switching to induction medium (DMEM/F12 containing 1 % Penicillin/Streptomycin (both from Invitrogen), 0.1 µM dexamethasone (Sigma-Aldrich), 100 nM insulin, 200 nM rosiglitazone (Sigma-Aldrich), 540 μM isobutylmethylxanthine (IBMX, Sigma-Aldrich), 2 nM T3 (Sigma-Aldrich) and 10 μg/ml transferrin (Sigma-Aldrich)). After 3 days of differentiation, IBMX was removed from the cell culture media. The cell cultures were left to differentiate for an additional 9 days.

Cells were fixed with 4% paraformaldehyde (PFA, 16% w/v ag. Soln., methanol free, Alfa Aesa) for 10 min at time points indicated in the figure and subsequently washed with PBS.

### Immunocytochemistry of cultured cells

Fixed cells were blocked with CT (0.5% Triton X-100 (Sigma-Aldrich) and 5% ChemiBLOCKER (Merck Millipore) in 0.1 M NaP, pH 7.0) for 30 min at room temperature. Primary antibody and secondary antibodies were diluted in CT and each incubated for 60 min at room temperature. As a DNA counterstain, DAPI was used (4’,6-Diamidino-2-Phenylindole, Dihydrochloride, 1:10,000, Invitrogen). For staining of lipid droplets, cells were incubated with the lipophilic dye LD540 (1:10,000) (Spandl *et al*., 2009) for 15 min and washed again with PBS. The following antibodies were used: rat anti-Ki-67 (1:500, Invitrogen, 14-5698-82), goat anti-Perilipin (1:400, Abcam, ab61682), donkey anti-rat-A647 (1:150, Dianova, 712-605-153), donkey anti-goat-Cy3 (1:1,000, Dianova, 705-165-147).

### Microscopy of cultured cells

Fluorescence images were taken at the CellDiscoverer7 widefield microscope (Zeiss) or the Observer.Z1 widefield microscope (Zeiss) using automated image acquisition. Four images were acquired per well, each in a z-stack (step size 4 µm, 10 x magnification). Depicted images are shown as a projection of the sharpest plane including the plane above and below. A maximum projection around the sharpest plane was generated using the ImageJ plugin ExtractSharpestPlane_JNH (https://doi.org/10.5281/zenodo.5646492) (Hansen, 2021).

### Mouse work

All animal experiments were performed in agreement with the German law of animal protection and local institutional animal care committees (Landesamt für Natur, Umwelt und Verbraucherschutz, LANUV). Mice were kept in individually ventilated cages in the mouse facility of University Hospital Bonn (Haus für Experimentelle Therapie (HET), Universitätsklinikum, Bonn). Mice were raised under a normal circadian light/dark cycle of each 12 h and animals were given water and complete- or very high-fat content (LARD) diet (ssniff Spezialdiäten) *ad libitum* (LANUV Az 81-02.04.2019.A170). At 11 weeks of age, a cohort of single-housed mice were switched to HFD for 7.5 weeks, while another was kept on CD. Before sacrifice using cervical dislocation, mice were anesthetized using isoflurane.

### Tissue fixation and histology

WAT was fixed for 24 h in 4% paraformaldehyde (PFA) at 4°C, before being further processed using the automated Epredia™ Excelsior™ AS Tissue Processor (Thermo Fisher Scientific™ Inc.). First, tissues were dehydrated by six incubation steps in increasing EtOH concentrations (70-100% at 30°C for 1 h each, UKB Pharmacy). This was followed by three steps in a clearing agent, xylene (30°C for 1 h each, AppliChem), to remove the ethanol, before incubating three times in molten paraffin wax (62°C for 80 mins each, Labomedic), which infiltrates the sample and replaces the xylene. Infiltrated tissues were then casted into molds together with liquid paraffin (65°C) and cooled to form a solid paraffin block with embedded tissue (Leica EG1150 H Paraffin Embedding Station and Leica EG1150 C Cold Plate). Paraffin-embedded WAT was sliced into 5 μm sections using a Thermo Scientific™ HM 355S Automatic Microtome and mounted on Surgipath® X-tra® Microscope Slides (Leica Biosystems). To represent the whole tissue, three different tissue depths were sliced and collected. Several 20 μm cutting steps were performed between each tissue depth.

WAT sections were stained for histological analysis with Mayer′s hemalum solution (Sigma Aldrich) and Eosin Y solution (1% in water, Roth) using the Leica ST5020 Multistainer combined with Leica CV5030 Fully Automated Glass Coverslipper. Deparaffinization of paraffin-embedded WAT slices was performed by two heating steps (60°C for 6 mins each) to melt the wax and three subsequent steps in xylene (1 min each), before incubation in a graded alcohol series (100%-70% ethanol; 80 sec each) to rehydrate the tissue sections and ending with a final rinsing step in sterile distilled water (dH2O) (80 sec). Next, tissue slices were stained with Mayer′s hemalum solution (3 mins), before washing in running tap water (5 mins). To counterstain with eosin, slides were immersed in eosin (25 sec) and then rinsed in dH_2_O (80 sec), before incubation on a graded alcohol series (70%-100% ethanol; 80 sec each) to dehydrate the tissue. After two final steps in xylene (60 sec each), stained slides were mounted with CV Mount (Leica Biosystems).

Paraffin embedding, slicing, and staining was conducted by the histology facility at University Hospital Bonn.

### Microscopy of tissues

Stained sections were stored at RT until imaging with the Zeiss Axio Scan.Z1 Slide Scanner at the Microscopy Core Facility of the Medical Faculty at the University of Bonn.

### Image analysis

For all presented data sets, we summarize the Image specifications, AdipoQ Preparator preferences, and AdipoQ preferences in Supplementary Table 2. Manual assessment of errors was performed in Image. Falsely detected objects were selected with the Wand tool in the mask to generate ROIs. These ROIs were then grouped by type of error (fused or fragmented), counted to determine the number of wrongly detected objects and the area was measured using ImageJ’s Measure function. Missed objects were quantified by manual counting in the image opened in ImageJ.

### Statistics

Statistical analysis was performed in GraphPad Prism (Version 9.3.0, GraphPad Software, Inc.). A Shapiro-Wilk test confirmed a normal distribution for all conditions in Figure 2B except HFD Female group 6000-7000 µm^2^ and CD Male groups 3000-4000 µm^2^ until >8000 µm^2^. However, these groups were confirmed to be normally distributed with a Shapiro-Wilk test when ignoring one largely outlying value. The standard deviation of compared conditions was very different for most comparisons (see bars in Figure 2B). Thus, unpaired, two-sided t-tests with Welch correction were applied.

### Hardware requirements

Any computer that can run ImageJ (Schneider *et al*., 2012; Rueden *et al*., 2017) or FIJI (Schindelin *et al*., 2012) is suited. ImageJ and FIJI run on Linux, Windows, and Mac operating systems.

### Code and software availability statement

The AdipoQ workflow involves two java-based ImageJ plugins that are open-source and freely available online through the GitHub repository: https://github.com/hansenjn/AdipoQ. The GitHub repositories provide the source code, the plugin files, a user guide, and allows to post issues or suggest new functions. The user guide describes all methods that are implemented into AdipoQ in detail.

### Software

Data analysis and statistical analysis was performed in Microsoft® Excel for Mac (Version 16.54), GraphPad Prism (Version 9.3.0, GraphPad Software, Inc.), R (R version 4.0.3 (2020-10-10)). The R Foundation for Statistical Computing), RStudio (Version 1.4.1103, RStudio, Inc.). All image processing and analysis was performed in ImageJ (Version 1.53a, U.S. National Institutes of Health, Bethesda, Maryland, USA). Plots and Figures were generated using GraphPad Prism (Version 9.3.0, GraphPad Software, Inc.), and Affinity Designer (Version 1.10.4, Serif (Europe), Ltd.). ImageJ plugins were developed in Java, with the aid of Eclipse IDE for Java Developers (Version 2021-09 (4.21.0), Eclipse Foundation, Inc., Ottawa, Ontario, Canada).

## Abbreviations

WAT: white adipose tissue
BAT: brown adipose tissue
AT: adipose tissue
HE: hematoxylin-eosin
HFD: high fat diet
CD: chow diet
APC: adipocyte progenitor cells
PLIN1: perilipin
Ads: adipocytes

## Acknowledgement

We would like to thank the Microscopy Core Facility of the Medical Faculty at the University of Bonn for providing help, services, and devices funded by the Deutsche Forschungsgemeinschaft (DFG, German Research Foundation) – project numbers 388168919, 388158066, 13123509; the Histology Platform of the ImmunoSensation^2^ Cluster of Excellence, Kim Dressler and E. Weidner for technical assistance, and Christoph Thiele (LIMES institute) for providing cells and the lipid dye LD540.

Research in the Wachten lab was supported by grants from the Deutsche Forschungsgemeinschaft (DFG) – SFB 1454 – project number 432325352, TRR83 – project number 112927078, SFB/TRR333, under Germany’s Excellence Strategy – EXC2151 – project number 390873048, as well as intramural funding from the University of Bonn. AP was supported by 335447717-SFB 1328 and 214362475-RTG1873/2. JNH was supported with a PhD fellowship from the Boehringer Ingelheim Fonds.

In Supplementary Figure S1, we show and analyze images from the Genotype-Tissue Expression (GTEx) Portal https://gtexportal.org/, retrieved on 05/18/2022. Accession numbers are: GTEX-15ER7-0426, GTEX-15EOM-0226, GTEX-12ZZX-0226, GTEX-1EN7A-0226, GTEX-1128S-2126, GTEX-11GSO-2326, GTEX-1PBJI-1326, GTEX-XQ3S-2426, GTEX-15SHU-2326, GTEX-13NYB-1126, GTEX-1269C-0726, GTEX-11DYG-2026. The Genotype-Tissue Expression (GTEx) Project was supported by the Common Fund of the Office of the Director of the National Institutes of Health, and by NCI, NHGRI, NHLBI, NIDA, NIMH, and NINDS.

## Author contribution statement

J.N.H, and D.W. conceived the project. K.S., N.W., M.H., P.L., D.R., T.G., A.P., D.W., and J.N.H. conceived and designed the experiments. K.S., N.W., M.H., P.L., D.R., and T.G. performed the experiments. K.S., N.W., M.H., P.L., D.R., and J.N.H. analyzed the data. J.N.H. designed and developed software. T.G. and A.P. provided methodology. K.S., N.W., D.W., and J.N.H. drafted the Article. K.S., N.W., and J.N.H. prepared the digital images.

