## Supplementary Information for "AdipoQ – a simple, open-source software to quantify adipocyte morphology and function in tissues and *in vitro*"

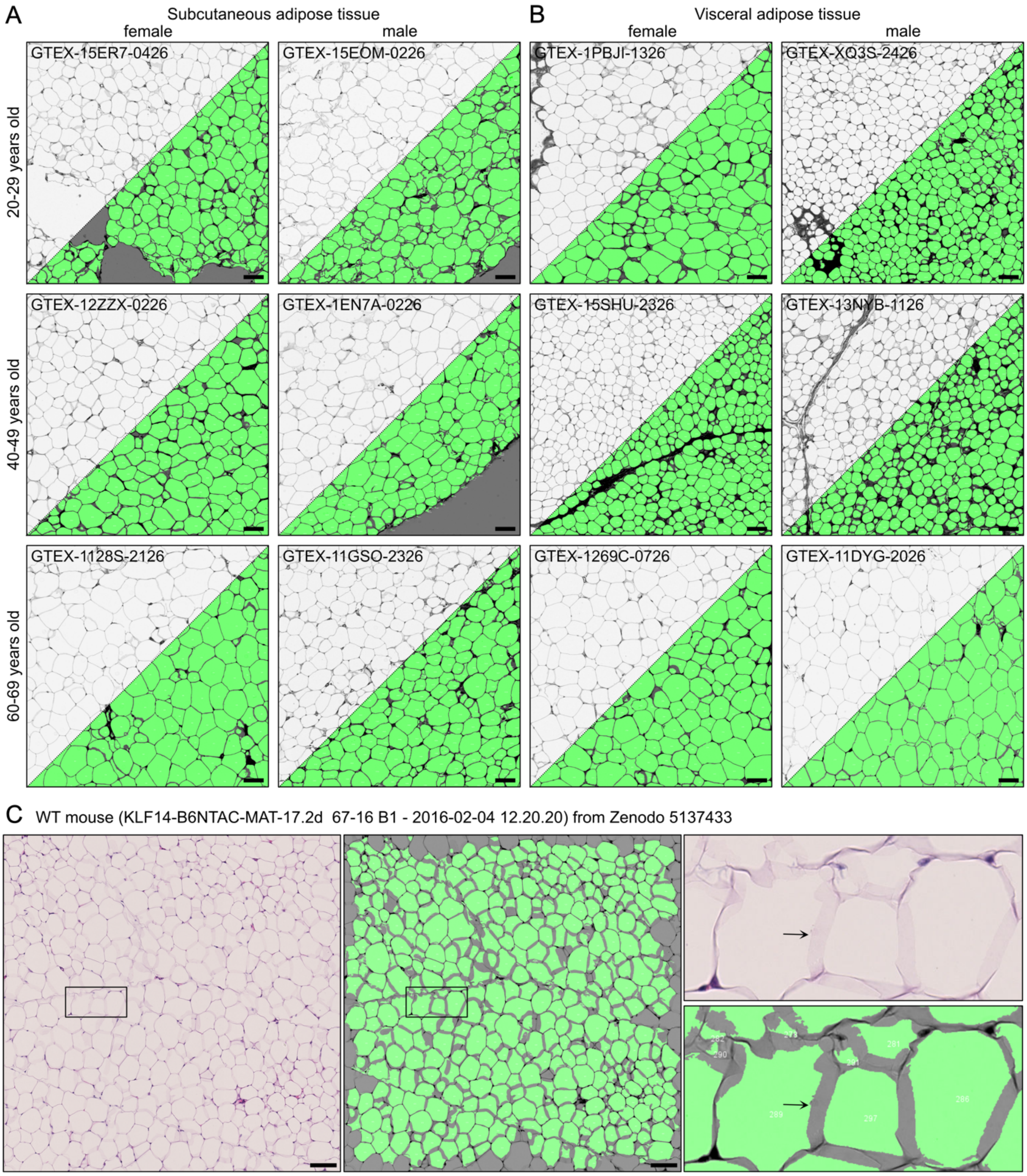


**Supplementary Figure 1: Analyzing images of adipose tissue sections from other labs with AdipoQ. (A, B)** Histological images of human visceral (A) and subcutaneous (B) adipose tissue from patients of different sex and age, downloaded from the Genotype-Tissue Expression (GTEx) portal (https://gtexportal.org/) on 05/18/2022. Accession numbers are indicated in the figure. The upper left shows the raw image, the lower right shows the raw image (reduced brightness) overlayed with the adipocyte masks detected by AdipoQ (green). (**C**) Image region extracted from a freely-available murine adipose tissue data set (Casero *et al.*, 2021) (https://dx.doi.org/10.5281/zenodo.5137433), published under a Creative Commons

Attribution 4.0 International license (https://creativecommons.org/licenses/by/4.0/legalcode). Left: raw image. Middle: raw image (reduced brightness) overlayed with the masks detected by AdipoQ (green). Right: Magnified views for the region marked on left. Arrows: Although borders between adipocytes are only weakly stained, AdipoQ detects individual adipocytes. Scale bars: 100 µm.

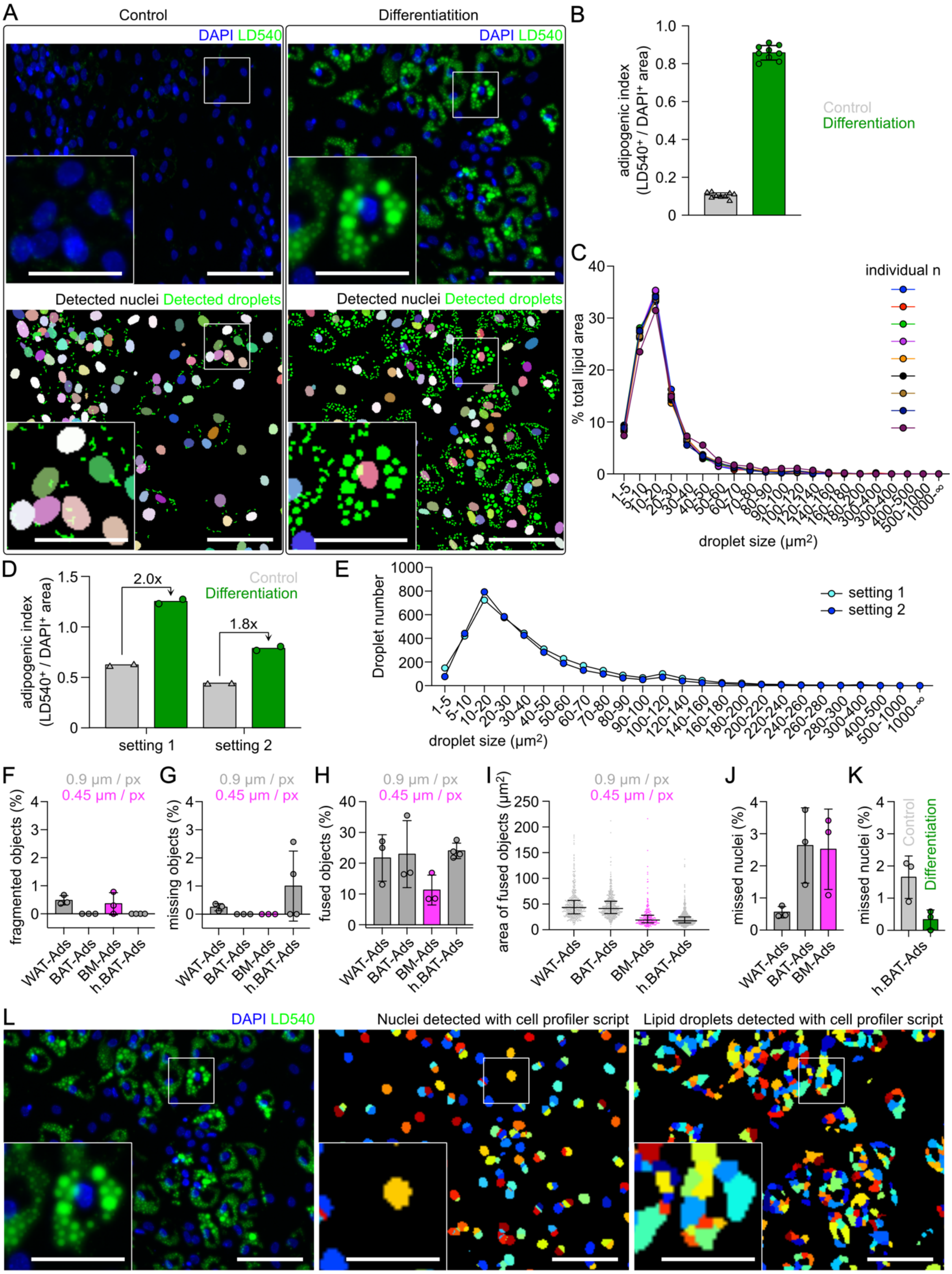


**Supplementary Figure 2: Analyzing adipocyte differentiation in a human, primary model for brown adipocytes. (A)** Example images from primary, human APCs after incubation with a control or differentiation medium. Rectangles indicate magnified views on lower left. Cells were stained with LD540 (green) and DAPI (blue). Nuclei were detected using the StarDist

ImageJ plugin (Schmidt *et al.*, 2018) (Model: “Versatile (fluorescent nuclei)”), implemented in AdipoQ. Each detected nucleus is shown in a different color. Droplets were detected using image processing and segmentation approaches implemented in AdipoQ. Top: raw fluorescence images. Bottom: detected nuclei and lipid droplets. Scale bars = 100 µm (magnified views: 50 µm). Exemplary images from n = 9 experiments (performed simultaneously). **(B-C)** Results from the AdipoQ analysis described in (A). **(B)** Adipogenic index, determined as the ratio of the total lipid droplet area ($LD540^+$) to the total nuclei area ($DAPI^+$). Bars show mean ± standard deviation. Data points show individual experiments. **(C)** Distribution of lipid droplet size, shown as the fraction of total lipid area, determined by AdipoQ analysis exemplified in (A). Lines and data points show individual experiments. **(D-E)** Results from an AdipoQ analysis with different AdipoQ Preparator settings (Setting 1 – Droplets: Blur image 0.9 µm, subtract blurred copy of the image 1.8 µm, custom threshold 20; Setting 2 – Droplets: Subtract background 18 µm, blur image 0.9 µm, subtract blurred copy of the image 1.8 µm, custom threshold 75; Setting 1 – Nuclei: Blur image 1.8 µm, subtract blurred copy of the image 2.7 µm, threshold algorithm Otsu; Setting 2 – Nuclei: StarDist prediction, default settings). Images shown for WAT-Ads in Figure 4A (day 7) and in Figure 4E were reanalyzed for this analysis. **(D)** Adipogenic index. Bars show mean. Data points show individual images. Fold changes of the mean adipogenic index for differentiated vs control cells are indicated at the arrows. **(E)** Distribution of lipid droplet size (absolute numbers per image, mean of both images shown). Lines and data points show individual analyses. **(F-K)** Manual assessment of errors in the object detection for randomly picked images whose results were shown in A-C and Figure 4E. For each image, a quarter of the image was analyzed. Individual data points show individual images (F-H, J-K) or individual detected objects (I). Bars indicate mean ± standard deviation (F-H, J-K) or median and interquartile range (I). Fragmented objects were defined as droplets detected as more than one object. Missing objects were defined as droplets (H) or nuclei (J-K) not detected. Fused objects were defined as detected objects corresponding to more than one droplet. Plotted is the percentage of objects from all detected objects (F-H, J-K). Gray and magenta data show images that featured a pixel width of 0.9 µm/px or 0.45 µm/px, respectively (F-J). h.BAT-Ads = human BAT-Ads. **(K)** Comparison of nuclei detection in differentiated and undifferentiated cultures. **(L)** Analysis of the image of differentiated human BAT-Ads shown in (A) with a CellProfiler script designed for lipid droplet detection (Adomshick *et al.*, 2020). Left: raw image, middle: detected nuclei, each detected nucleus is shown in a different color, right: detected droplets, each detected droplet is shown in a different color. Bars and magnified views as described for (A).

**Supplementary Table 1: AdipoQ output parameters.** Intensity parameters are determined and output separately for any channel in the image. Parameters starting with “Surr” are referring to the pixels in the surroundings of the object. These pixels are defined as the pixels that do not belong to the object but whose Euclidian distance to at least one pixel belonging to the object is less than a user defined value (demonstrated in Figure 2E).

| **Parameter label** | **Units** | **Description** |
|---|---|---|
| Center X, Y, Z | µm | Center of the object. Indicates the position in the image. |
| Voxels | - | The number of pixels that belong to the object. Holes in the object are not included. |
| Area | $\mu m^2$ | The area of the object, calculated as the product of the number of pixels, the pixel width, and the pixel height. |
| Outline | µm | The pixelated outline of the object, determined as the sum of the individual pixel outlines with each pixel’s outline defined as the border outlines of the pixel, at which no neighbored pixels are present that belong to the object. |
| 2D-Asphericity Index | - | Ratio of the outline of the object to the outline of a perfect circle containing the same area as the object. This parameter gives a measure on how aspherical the object’s shape is. Determined as $outline/(\sqrt{\pi \cdot area})$. An index of 1 represents a perfectly circular shape. The higher the index, the less circular the shape.<br>This parameter could be used in a post-hoc analysis to detect and remove objects that do not show an adipocyte-, droplet- or nucleus-like shape. |
| Average Intensity | - | Average of the intensity values of all pixels belonging to the object. |
| Integrated Intensity | - | The sum of the intensity values of all pixels belonging to the object. |
| Median Intensity | - | Median of the intensity values of all pixels belonging to the object. |
| SD of Intensities | - | Standard deviation of the intensity values of all pixels belonging to the object. |
| Min Intensity | - | Lowest intensity value among all pixels belonging to the object. |
| Max Intensity | - | Highest intensity value among all pixels belonging to the object. |
| Surr Voxels | - | The number of pixels that are considered to belong to the surroundings. |
| Surr Average Intensity | - | Average of the intensity values of pixels in the surroundings. |

| | | |
|---|---|---|
| Surr Integrated Intensity | - | The sum of the intensity values of pixels in the surroundings. |
| Surr Median Intensity | - | Median of the intensity values of pixels in the surroundings. |
| Surr SD of Intensities | - | Standard deviation of the intensity values of pixels in the surroundings. |
| Surr Min Intensity | - | Lowest intensity value among all pixels in the surroundings. |
| Surr Max Intensity | - | Highest intensity value among all pixels in the surroundings. |
| Surr Average Intensity Min 5% | - | Average of the 5% lowest intensity values among all pixels in the surroundings. |
| Surr Average Intensity Min 25% | - | Average of the 25% lowest intensity values among all pixels in the surroundings. |
| Surr Average Intensity Max 5% | - | Average of the 5% highest intensity values among all pixels in the surroundings. |
| Surr Average Intensity Max 25% | - | Average of the 25% highest intensity values among all pixels in the surroundings. |

**Supplementary Table 2: Image preferences and AdipoQ settings for presented data sets.** Hyphen (-) denotes that the respective function was not applied or the setting was irrelevant. Custom threshold values were calculated based on the background intensity level, see the User Guide on the GitHub page for details.

| | **Fig. 2**<br>**Suppl. Fig. 1** | **Fig. 3** | **Fig. 4A** | **Fig. 4B** | **Fig. 4C** | **Suppl. Fig. 2** |
|---|---|---|---|---|---|---|
| **Microscopy and input image specifications** | | | | | | |
| Microscope | Zeiss Axio Scan.Z1 | Zeiss Observer Z.1 | Zeiss CD7 | Zeiss CD7 | Zeiss CD7 | Zeiss CD7 |
| Channel 1 | Red | DAPI | Ki-67 | DAPI | DAPI | LD540 |
| Channel 2 | Green | PLIN1 | DAPI | LD540 | LD540 | DAPI |
| Channel 3 | Blue | Ki-67 | LD540 | Transmit-ted light | Transmit-ted light | Transmit-ted light |
| Channel 4 | - | Transmitted light | Transmitted light | - | - | |
| Objective | 10x | 10X | 20X, Afocal magnification changer 0.5X | | | |
| Image size | Diverse | 2752x2208 | 980x708 | 980x708 | 1956x1416 | 980x708 |
| Pixel size (µm) | 0.442 | 0.227 | 0.903 | 0.903 | 0.452 | 0.910 |
| **AdipoQ Preparator** | | | | | | |
| **Channel to be segmented #1** | | | | | | |
| Channel Nr | 2 | 1 | 2 | 1 | 1 | 1 |
| Subtract background - radius (µm) | Fig. 2A-C, Suppl. Fig. 1C: -<br>Fig. 2E-F: 22.1 µm<br>Suppl. Fig. 1A, B: 30 µm | - | - | - | - | - |
| Blur - sigma (µm) | - | 8 | - | 1.820136 | - | 0.455034 |
| Subtract blurred copy - sigma (µm) | - | 10 | - | 2.730204 | - | 0.910068 |
| Segmentation method | Triangle | Li | StarDist | Custom: 40 | StarDist | Custom: 50 |
| Background definition | Dark on bright background | Bright on dark background | | | | |
| Exclude zero pixels - gap radius | 2.2 | - | - | - | - | - |

| | | | | | | |
|---|---|---|---|---|---|---|
| (µm) | | | | | | |
| Despeckle | Yes | - | - | - | - | - |
| Detect tissue regions - minimum radius (µm) | Yes, 8.8 | - | - | - | - | - |
| During detection, close gaps | - | - | - | - | - | - |
| Fill holes | Yes | Yes | Yes | Yes | Yes | Yes |
| Watershed | Fig. 2: Yes<br>Suppl. Fig. 1: No | Yes | Yes | Yes | Yes | Yes |
| **Channel to be segmented #2** | | | | | | |
| Channel Nr | - | 3 | 3 | 2 | 2 | 2 |
| Subtract background - radius (µm) | - | - | 18.059131 | - | - | - |
| Blur - sigma (µm) | - | 3 | 0.902957 | 0.9100680 | 0.677217 | - |
| Subtract blurred copy - sigma (µm) | - | 6 | 1.805913 | 1.820136 | 1.354434 | - |
| Segmentation method | - | Custom: 50 | Custom: 75 | Custom: 20 | Custom: 20 | StarDist |
| Background definition | - | Bright on dark background | | | | |
| Exclude zero pixels - gap radius | - | - | - | - | - | - |
| Despeckle | - | - | - | - | - | - |
| Detect tissue regions - minimum radius | - | - | - | - | - | - |
| During detection, close gaps | - | - | - | - | - | - |
| Fill holes | - | Yes | Yes | Yes | Yes | - |
| Watershed | - | Yes | Yes | Yes | Yes | - |
| **AdipoQ Analyzer** | | | | | | |
| Channel Nr | 2 | 3 | 2&4 | 1&3 | 1&3 | 1 & 3 |
| Increase range | Yes | - | - | - | - | - |
| Minimum particle size (µm$^2$) | 20 | 1.03058 | Channel 1: 18.059131<br>Channel 2: 4.514783 | Channel 1: 18.20136<br>Channel 2: 4.55 | Channel 1: 20<br>Channel 2: 5 | Channel 1: 4.55034<br>Channel 3: 8.282237 |

| | | | | | | |
|---|---|---|---|---|---|---|
| Additionally exclude | Particles touching x or y borders | Nothing | Nothing | Nothing | Nothing | Nothing |
| Quantify surrounding (µm) | Yes, 8.84 | - | - | - | - | - |
| Fuse included particles | - | - | - | - | - | - |
